# Hi-C data from the filamentous fungus *Podospora anserina* and associated 3D models to visualize the spatial organization of its chromosomes

**DOI:** 10.64898/2026.06.14.732207

**Authors:** Goran Royer, Anakim Gualdoni, Pierre Poulain, Fabien Dumetz, Nadia Ponts, Pierre Grognet, Fabienne Malagnac, Gaëlle Lelandais

## Abstract

**Objectives:** Model species are essential for fundamental research in biology. While a complete genomic sequence is a prerequisite for genetic studies, it is not enough on its own. Understanding the three-dimensional organization of the genome is also important, allowing researchers to gain a more realistic understanding of the mechanisms governing genome function. In the fungal model *Podospora anserina*, although the genomic sequence has been established for a long time, the three-dimensional organization remained unknown. Here we obtained the first Hi-C datasets and present associated 3D models, providing the research community with a valuable resource for better multi-omics data integration.

**Data description:** Hi-C experiments were performed in duplicate, using nuclei purified from wild-type fungal mycelium. Four FASTQ files were obtained (two per replicate) and used as inputs for the 3DGB workflow with four different output resolutions, to observe the genome organization of *P. anserina* at different levels of detail (50 kb, 20 kb, 10 kb, and 5 kb). In a context where researchers already have, for this species, a large amount of traditional omics data (ChIP-seq, RNA-seq, etc.), these 3D models are helpful for complementing the linear representation of the genome, which is traditionally used in bioinformatic analyses.

## Objective

In our laboratory, we study the species *Podospora anserina*, an ascomycete fungus that is used as a model system for the study of several biological processes such as sexual reproduction, cell differentiation and death, mitochondrial physiology or ageing [1]. Since its genome was completely sequenced and annotated [2, 3], multiple functional “omics” studies have been conducted on this species [2, 4–7]. The question of integrating this data, beyond simply comparing lists of genes or superimposing tracks using genome visualization tools such as IGV [8] or ARTEMIS [9] is a new challenge for current functional genomics projects.

By integrating a crucial regulatory layer for gene expression, 3D models of genome organization provide an original perspective for re-evaluating both existing and emerging functional data. In a previous study, we introduced 3D models of chromosomes from several fungal species [10]. These models were obtained using a bioinformatics workflow called “3D genome builder” (3DGB). Open source and freely available on GitHub (https://github.com/data-fun/3d-genome-builder), this workflow has the advantage of linking the creation of contact maps with the building of 3D models. It also adds further processing of the 3D model output as PDB files, suitable for advanced visualization with molecular viewer software, and hence it helps to perform visual integrations of omics data (ChIP-seq, RNA-seq, etc.). 3DGB offers a comprehensive view of genomes that reinforces intuition and establishes the basis for developing new ideas [10].

Therefore, knowledge of the 3D organization of a genome is valuable information, helping to better understand the regulatory mechanisms which underly genome function [11]. In this article, we present the first 3D models of the *P. anserina* genome. These were obtained from an original, high-quality and reproducible Hi-C dataset, enabling us to examine the genome’s overall organization at various resolutions, ranging from 50 kb to 5 kb.

## Data description

Hi-C experiments were performed in duplicate using purified nuclei from wild-type mycelium. Four FASTQ files were thus obtained (two per replicate) and are shared in **DataSet 1** (see **Table 1**). The quality of the sequences was checked to ensure robust model construction using 3DGB (**DataFile 1**).

**Table 1:** Overview of data files/data sets.

| Label | Name of DataFile /data set | File types (file extension) | Data repository and identifier (DOI or accession number) |
| --- | --- | --- | --- |
| DataFile 1 | General statistics of the FASTQ files (quality scores and output statistics from the Hi-Pro software) | Portable Document Format file (.pdf) | Zenodo: https://doi.org/10.5281/zenodo.22039028 [14] |
| DataFile 2 | Configuration file to run the 3DGB workflow | Portable Document Format file (.pdf) | Zenodo: https://doi.org/10.5281/zenodo.22039295 [15] |
| DataFile 3 | Images of contact maps obtained at different resolutions and associated 3D models | Portable Document Format file (.pdf) | Zenodo: https://doi.org/10.5281/zenodo.22039558 [16] |
| DataFile 4 | Explanation of the interpolation procedure used to create a 3D model with gene locations | Portable Document Format file (.pdf) | Zenodo: https://doi.org/10.5281/zenodo.22039736 [17] |
| DataFile 5 | Script to calculate 3D model 'per gene', from 3D model 'per bin' | Python script (.py) | Zenodo: https://doi.org/10.5281/zenodo.22040019 [18] |
| DataFile 6 | 3D coordinates for beads associated with "genes" at 10kb resolution | Protein Data Bank (.pdb) | Zenodo: https://doi.org/10.5281/zenodo.22040162 [19] |
| DataFile 7 | Example of visual integration of ChIP-seq data onto the 3D model "per bin" at 10 kb resolution | Portable Document Format file (.pdf) | Zenodo: https://doi.org/10.5281/zenodo.22040206 [20] |
| DataFile 8 | Example of a gene family representation onto the 3D model "per gene" at 10kb resolution | Portable Document Format file (.pdf) | Zenodo: https://doi.org/10.5281/zenodo.22040230 [21] |
| DataSet 1 | Raw FASTQ files obtained from HiC experiments | FASTQ file (.fastq) | ENA: http://identifiers.org/bioproject:PRJEB114401 [22] |
| DataSet 2 | Contact matrices at different resolutions used to create the 3D models with 3DGB | Portable Network Graphics (.png) | Zenodo: https://doi.org/10.5281/zenodo.22040269 [23] |
| DataSet 3 | 3D coordinates for beads associated with 'bin' at different resolutions | Protein Data Bank file (.pdb) | Zenodo: https://doi.org/10.5281/zenodo.22040312 [24] |

The 3DGB parameter file is presented in **DataFile 2**. In addition to the FASTQ files, it is necessary to provide the reference genome sequence, information regarding the restriction enzymes used in the Hi-C experiment, and the desired output resolution. In this work, we created models for four different resolutions (50 kb, 20 kb, 10 kb, and 5 kb), thus allowing visualization of the *P. anserina* chromosome organization with varying levels of detail. The 3DGB output files are shared across **DataFile 1** (analysis exploration graphs), **DataSet 2** (text files of contact frequencies), and **DataSet 3** (PDB files of 3D coordinates of interacting genomic regions). Note that images of the contact frequencies, along with a visualization of the associated 3D models, are shown in **DataFile 3** for each resolution. Notably, the models obtained at different resolutions shared a common organization that was consistent with the classical “Rabl” conformation, *i*.*e*., a particular positioning of chromosomes in which their centromeres and telomeres are respectively clustered at different peripheral locations, inside the nuclear envelope [10].

By default, 3DGB calculates models “by intervals” or “bins”, as defined in Hi-C data analysis protocols [12]. To go further in these representations, we wrote a Python script that interpolates, from the “bin model”, relevant x, y and z coordinates for a list of any genomic elements, for which locations on the original genomic sequence are known. The explanation of how the procedure works is provided in **DataFile 4** and the associated script is shared in **DataFile 5**. With this script, we calculated a “by gene” model (**DataFile 6**).

Examples of how the 3D models of the *P. anserina* genome can be used for visual integration of omics data are provided in **DataFile 7** (10kb model “per bin”) and **DataFile 8** (10kb model “per gene”). In **DataFile 7**, we present genome-wide patterns of three post-translational histone modifications, relating to open euchromatin (H3K4me3) and compact heterochromatin (H3K9me3 and H3K27me3), as described in the reference [7]. As expected, we observed that heterochromatin marks H3K9me3 and H3K27me3 are located at the periphery of the nuclear compartment, which is different from the location of the euchromatin mark H3K4me3. Such an observation is relevant, given the very different functional roles of these two chromatin states (respectively referred to as “closed” and “open” chromatin). In **DataFile 8**, we show the relative locations of genes that belong to biosynthetic gene cluster (BGC) [4]. BGC is a typical arrangement of genes that encode enzymes belonging to the same metabolic pathway (for example a toxin or antibiotic) and often sit in dynamic regions of chromosomes (subtelomeric regions or areas rich in mobile elements) [13]. In this study, we observed interesting inter-cluster groupings, particularly on chromosome 5, where the 3D structure is most constrained.

## Limitations

With this article, our goal is to make freely available and document the 3D models of the *P. anserina* genome, a model species for several laboratories worldwide. Although the underlying dataset is small (two replicates for one strain, studied at one development stage), which represents a notable limitation, it represents the first Hi-C data ever obtained for this species, making it a unique and valuable resource. We have made important efforts to follow best practices for data and code sharing, allowing anyone to easily reproduce our work. Like a complete reference genome sequence, these models provide essential information for better understanding the functioning of the *P. anserina* genome. Given its position as a model species in the fungal kingdom, sharing this data offers an interesting opportunity for the research community to review previous studies or initiate new projects that account for the structural properties of the genome.

## Supporting information

DataFile1

DataFile2

DataFile3

DataFile4

DataFile5

DataFile6

DataFile7

DataFile8

## Declarations

### Ethics approval and consent to participate

Not applicable.

### Consent for publication

Not applicable.

### Availability of data and materials

See **Table 1** for details and links to the data. Sequencing data have been deposited in the ENA under accession PRJEB114401 [22], result files are available on Zenodo [14–21, 23, 24], and the analysis codes are hosted on the CNRS GitLab (https://src.koda.cnrs.fr/AnakimGualdoni/3dgb-chipseq-analysis) [25].

### Authors’ contributions

Preparation of biological samples: P.G. and F.D.; Analysis of sequencing data: G.R. and G.L.; Creation of 3D models: G.R, P.P., A.G., and G.L.; Data integration: A.G. and G.L.; Study design: N.P., F.M; Writing: G.L; Review and editing: P.G, P.P, N.P, and F.M.

## Acknowledgements

Not applicable.

