## Supplementary figures and images for "Hi-C data from the filamentous fungus *Podospora anserina* and associated 3D models to visualize the spatial organization of its chromosomes"

### contact_map_5000.png

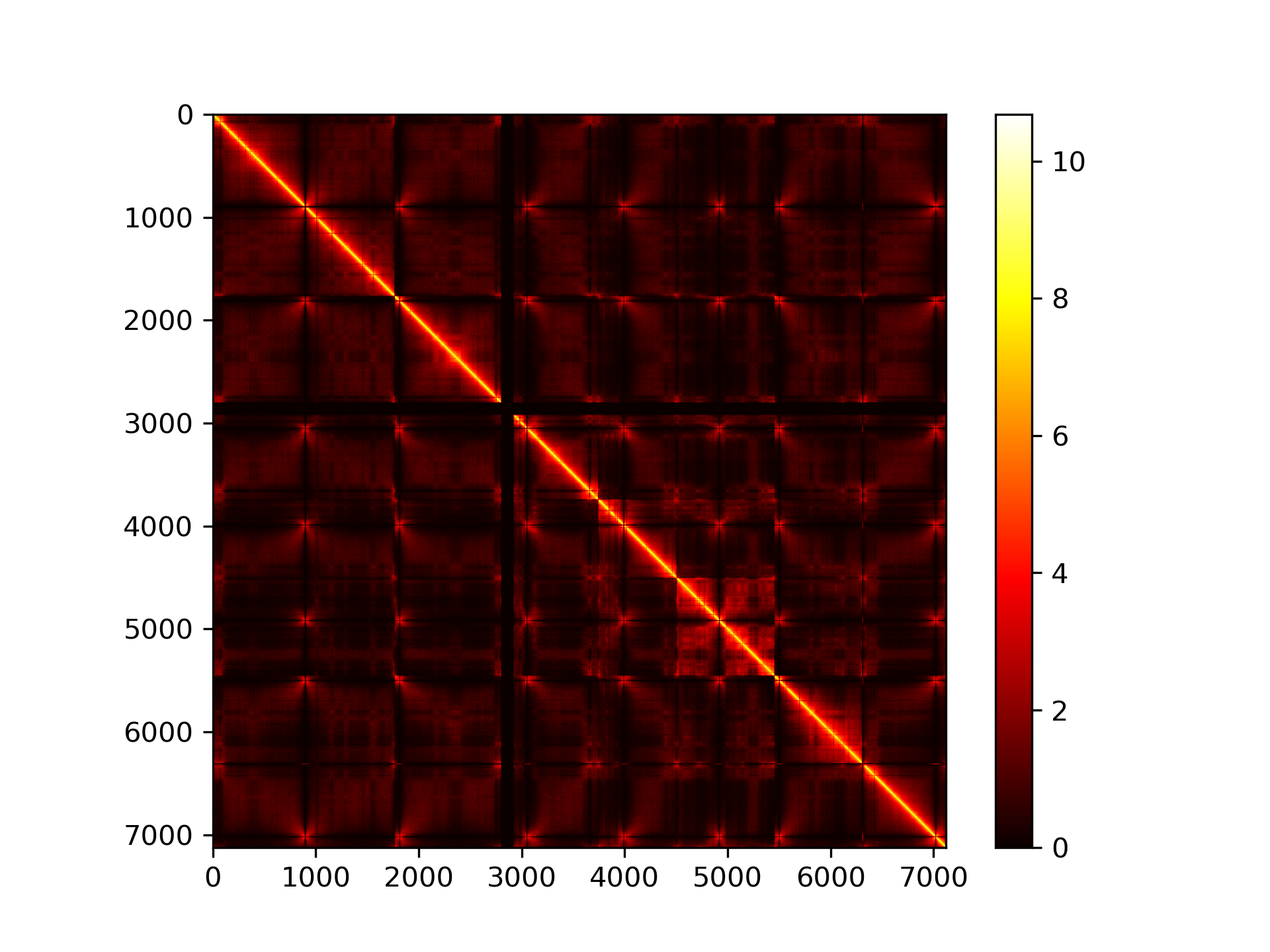

### contact_map_10000.png

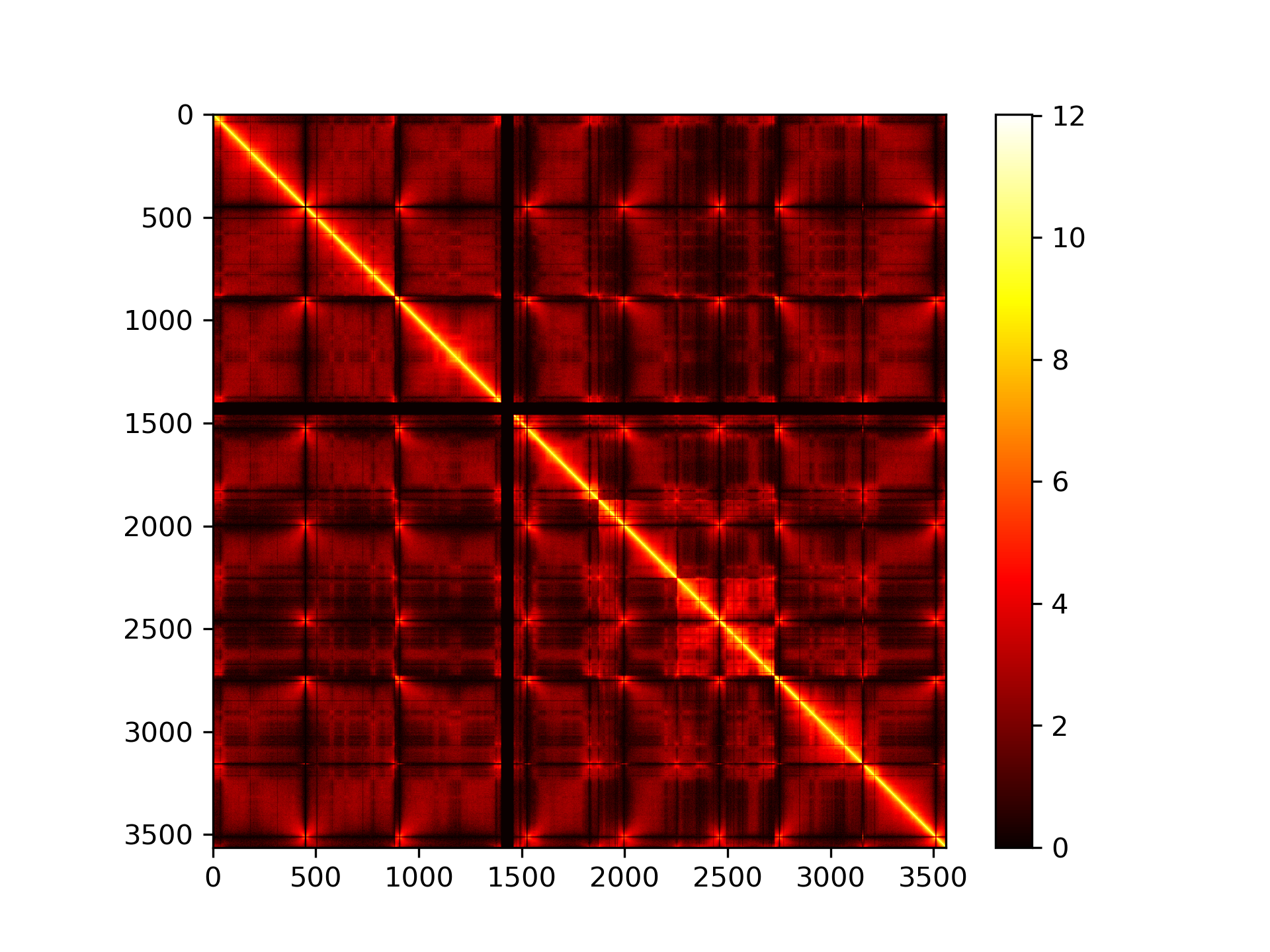

### contact_map_20000.png

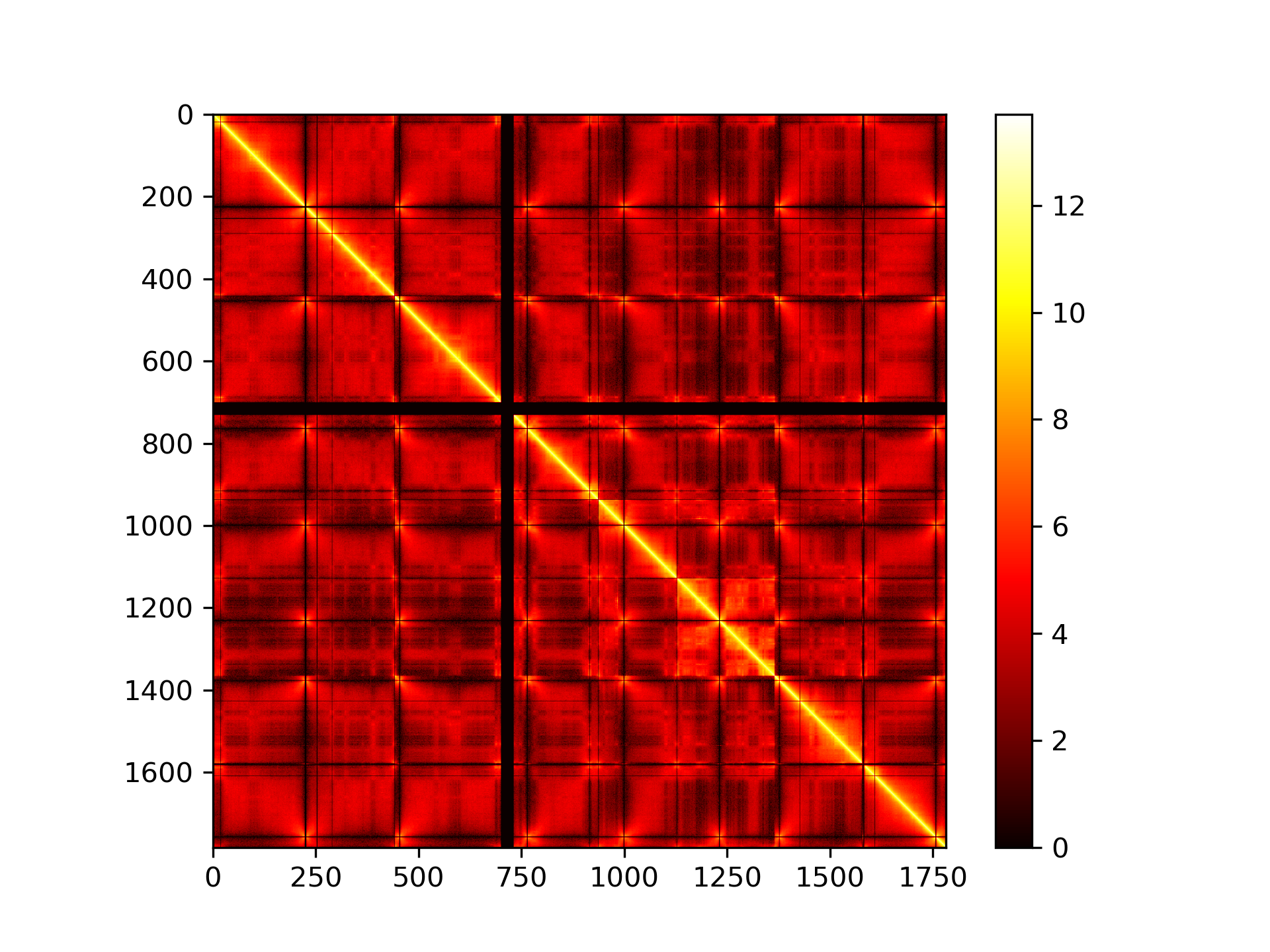

### contact_map_50000.png

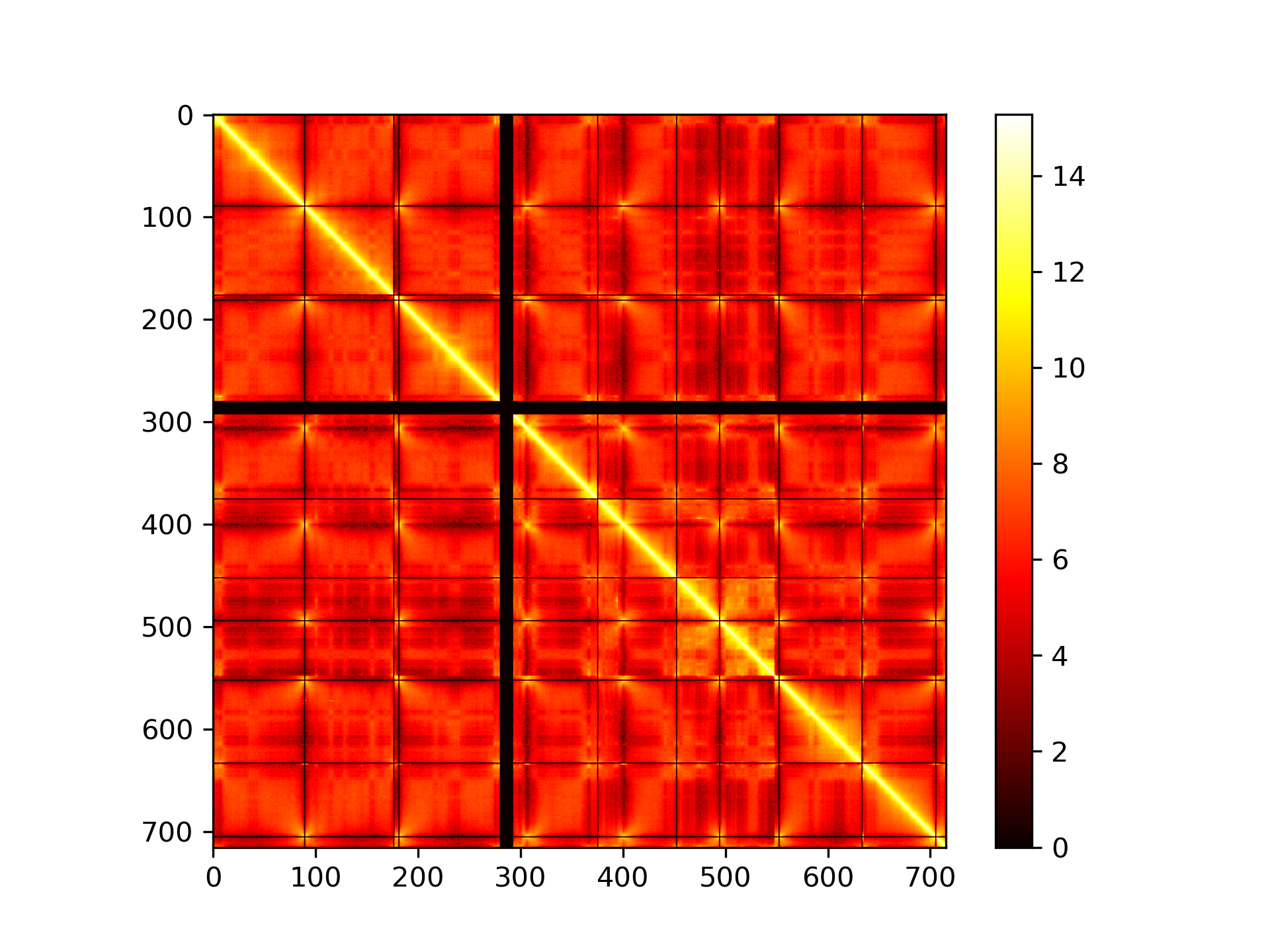
