## Supplementary material for "Hi-C data from the filamentous fungus *Podospora anserina* and associated 3D models to visualize the spatial organization of its chromosomes": DataFile1

### DataFile 1

Detailed information concerning the raw data files (FASTQ) used to create 3D models of the *Podospora anserina* genome using 3DGB software (1) (**Table S1.1**, **Figure S1.1** and **Table S1.2**), and output statistics from the Hi-Pro software (2) regarding the analysis of FASTAQ files with 3DGB (1) (**Figure S1.2** and **Figure S1.3**).

| File name | Number of reads | Initial read length | % duplicated reads | % GC |
|---|---|---|---|---|
| **WT1_R1.fastq** | 71.3 millions | 150 bp | 48.3 | 51 |
| **WT1_R2.fastq** | 71.3 millions | 150 bp | 41.0 | 52 |
| **WT2_R1.fastq** | 51.7 millions | 150 bp | 42.6 | 51 |
| **WT2_R2.fastq** | 51.7 millions | 150 bp | 35.9 | 52 |

**Table S1.1 : General statistics of the FASTQ files obtained from the MultiQC program (3).**

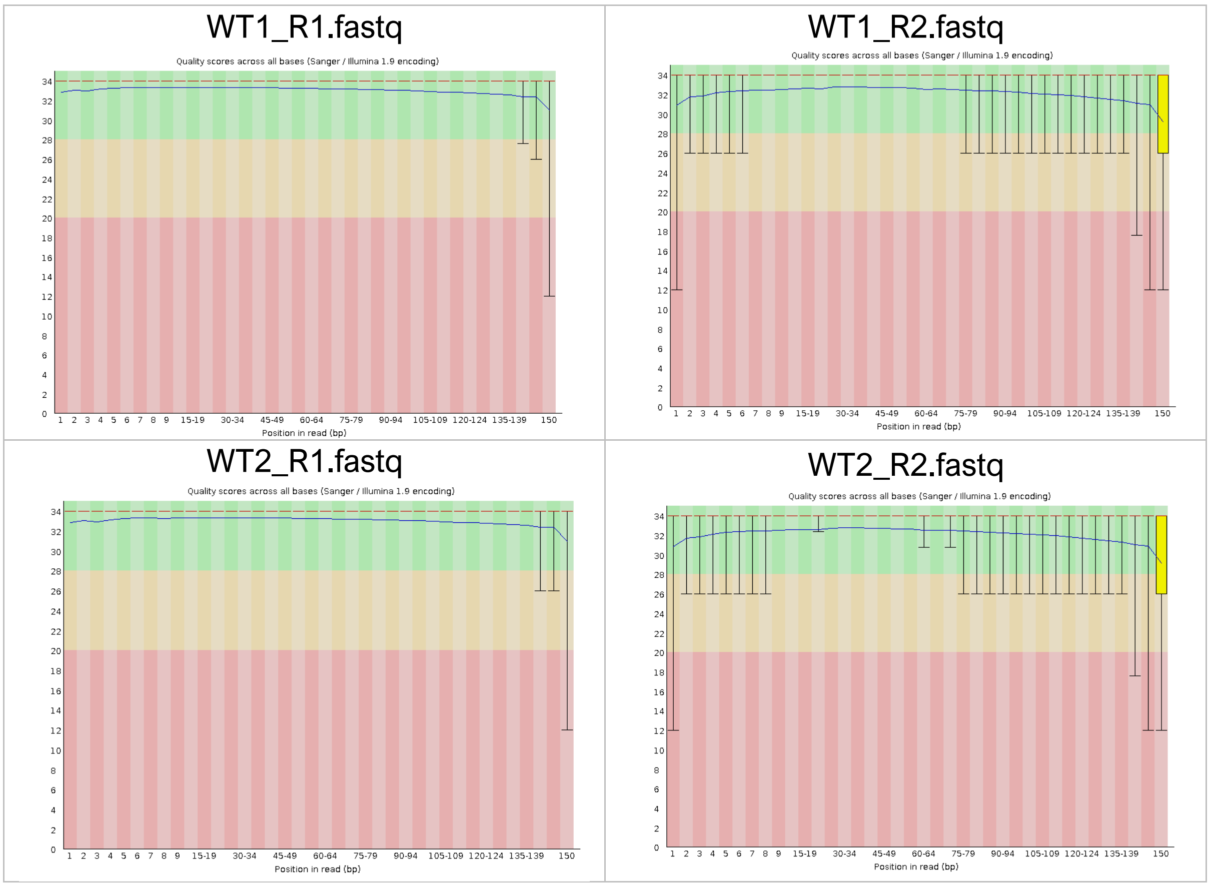


**Figure S1: Graphical representations of the quality scores across all bases in each FASTQ file, obtained from the FASTQC program** https://www.bioinformatics.babraham.ac.uk/projects/fastqc/**.**

| | Aligned reads (R1) | Aligned reads (R2) | Reported pairs |
|---|---|---|---|
| WT1 replicate | 89% | 85.4% | 68.7% (~50 millions) |
| WT2 replicate | 88.7% | 85.1% | 68.4% (~35 millions) |

**Table S1.2: General statistics of the FASTQ file analyses, performed with HiC-Pro (2).**

- <u>WT1 replicate</u>

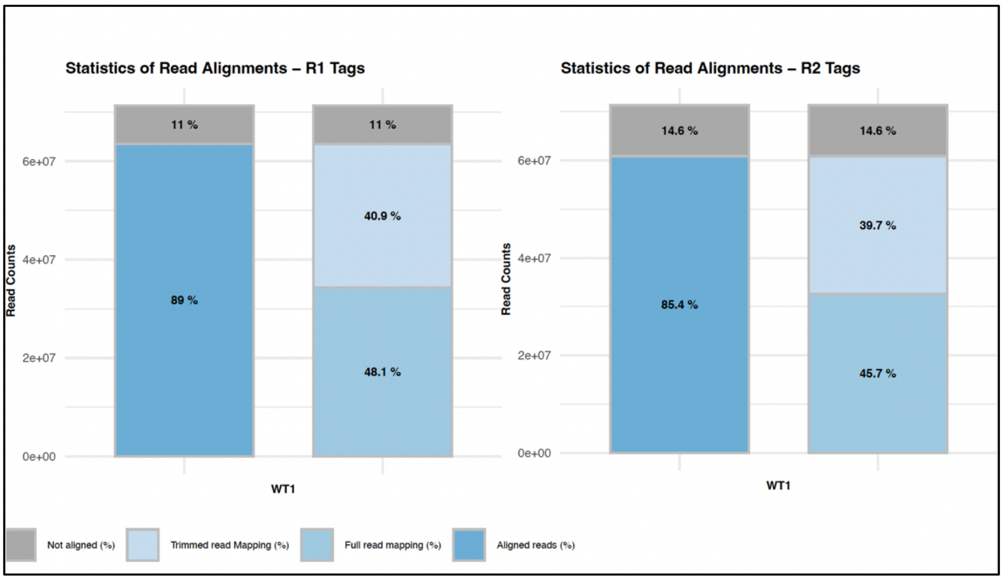


- <u>WT2 replicate</u>

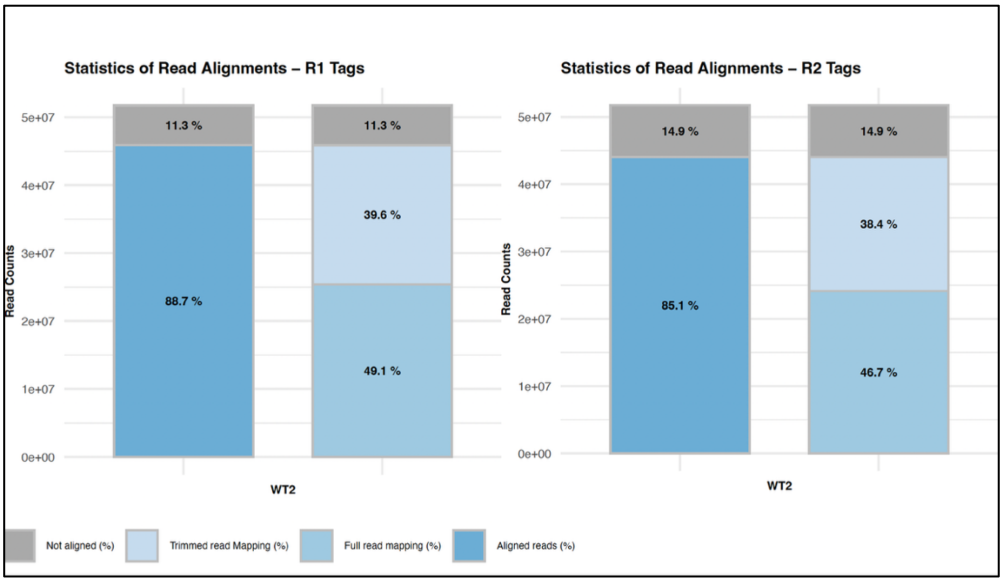


**Figure S1.2: Graphical representations of statistics from the Hi-C Pro software, associated with the mapping step. Detailed graphic explanations can be found here:** https://nservant.github.io/HiC-Pro/RESULTS.html#mapping-results.

- WT1 replicate

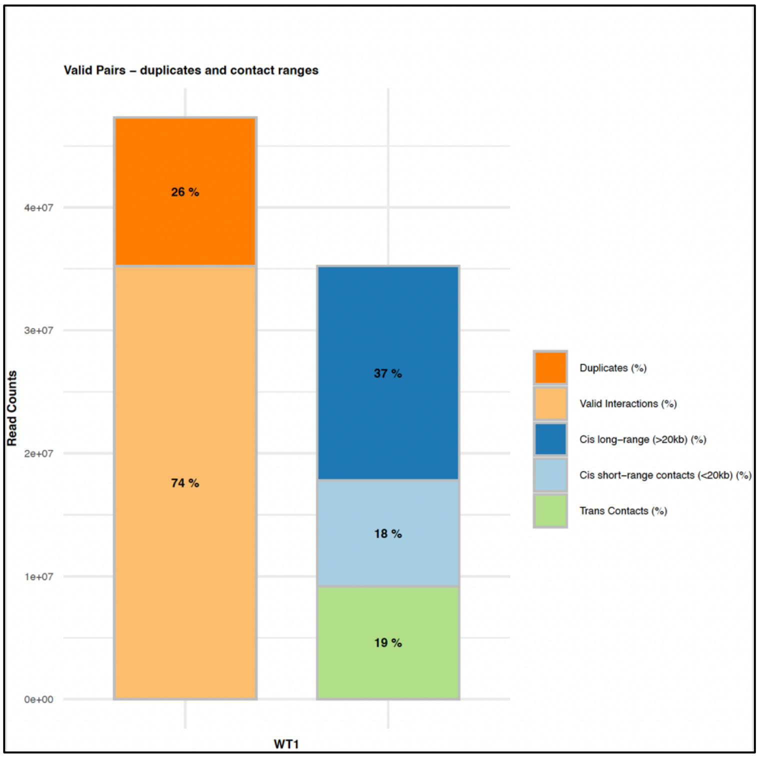


- WT2 replicate

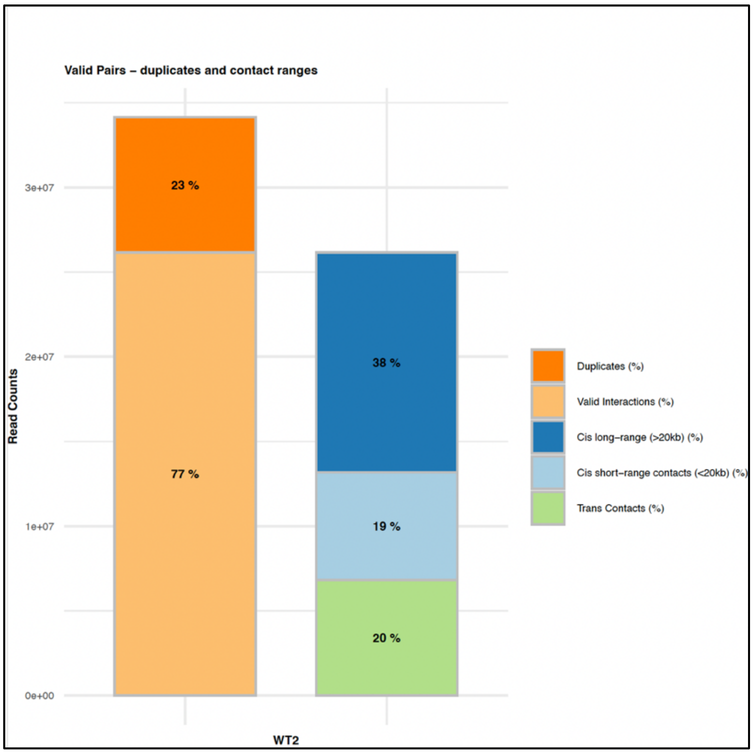


**Figure S1.3: Intra- and inter-chromosomal contact maps identified using Hi-C Pro. Detailed graphic explanations can be found here:** [https://nservant.github.io/HiC-Pro/RESULTS.html#intra-and-inter-chromosomal-contact-maps](https://nservant.github.io/HiC-Pro/RESULTS.html#intra-and-inter-chromosomal-contact-maps).
