## Supplementary material for "Hi-C data from the filamentous fungus *Podospora anserina* and associated 3D models to visualize the spatial organization of its chromosomes": DataFile2

### DataFile 2

Configuration file used to run the 3DGB workflow, following the documentation available at https://github.com/data-fun/3d-genome-builder?tab=readme-ov-file#create-the-config-file. Four different resolutions are specified to obtain 3D models with increasing levels of detail.

p_anserina_WT_3D-paper.yml — YAML

verify_contigs:

```yaml
# Wording directory name, without space
workdir: "3DGB_p_anserina_WT_3D-paper"

# Organism name
organism: "podospora_anserina"

# Sample identifiers
sra_ids:
- WT1
- WT2

# Restriction sites
hicpro_restriction_sites: "^GATC,G^ANTC"

# Ligation site
# This parameter is optional.
# And can contain multiple ligation motifs separated by comma.
hicpro_ligation_site: "GATCGATC,GANTGATC,GANTANTC,GATCANTC"

# Output resolutions for HiC-Pro
hicpro_resolutions:
- 50000
- 20000
- 10000
- 5000

# Output resolutions for Pastis
# should match the resolutions used in HiC-Pro.
# Fewer resolutions can be used.
pastis_resolutions:
- 50000
- 20000
- 10000
- 5000

# Verify inverted contigs
# Could be True or False
verify_contigs: False

```

Lignes : 39 Caractères : 726 Position : 726 Ligne : 39 726 octets Unicode (UTF-8) LF
