## Supplementary material for "Hi-C data from the filamentous fungus *Podospora anserina* and associated 3D models to visualize the spatial organization of its chromosomes": DataFile3

### DataFile 3

Images of contact frequency maps (on the left) and associated 3D models obtained for different resolutions (on the right). The contact maps were visualized using 3DGB software (1), and the 3D models were visualized using Mol* software (2). Each analysis was performed independently, using the FASTQ files presented in **Data File 1** and the 3DGB configuration file presented in **Data File 2**. These models correspond to what are known as 'bin' models, meaning that each bead represents a genomic region of a constant size (50kb, 20kb, 10kb or 5kb). The number of beads in each model is indicated in the last table.

- 50 kb:

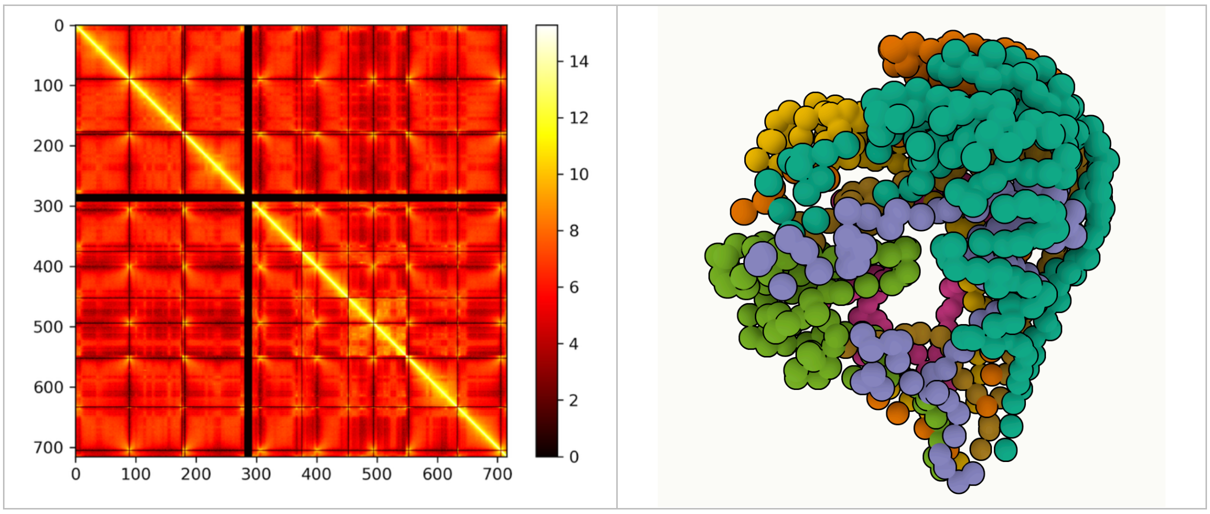


- 20 kb:

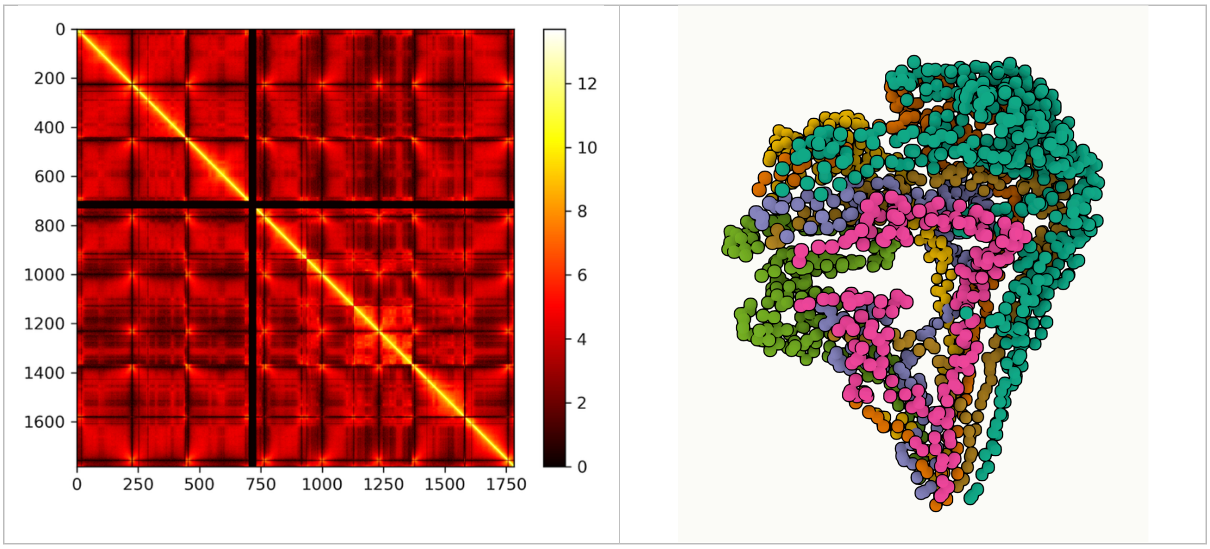

- 10 kb:

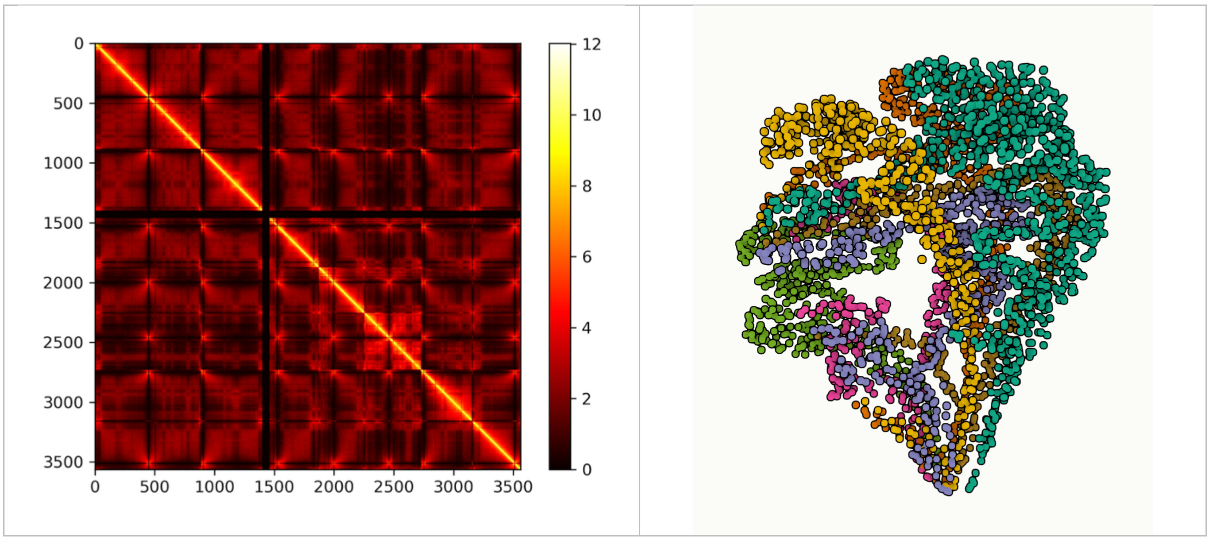


- 5 kb:

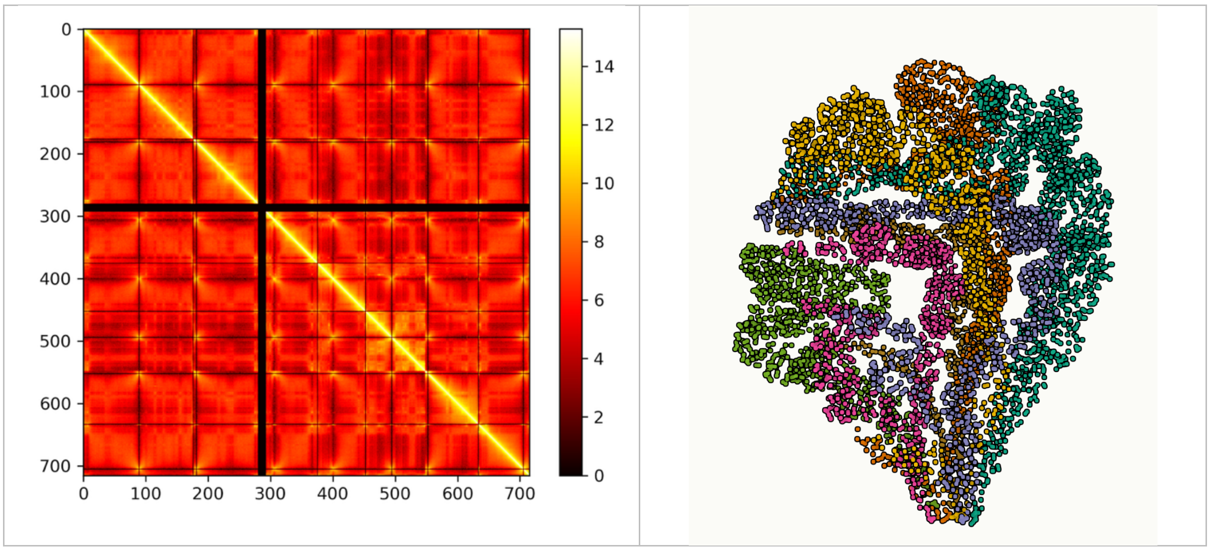


| BIN SIZE | # OF BEADS IN THE 3D MODEL (CLEANED) |
|---|---|
| **50 KB** | 683 |
| **20 KB** | 1 666 |
| **10 KB** | 3 238 |
| **5 KB** | 6 527 |
