## Supplementary material for "Hi-C data from the filamentous fungus *Podospora anserina* and associated 3D models to visualize the spatial organization of its chromosomes": DataFile4

### DataFile 4

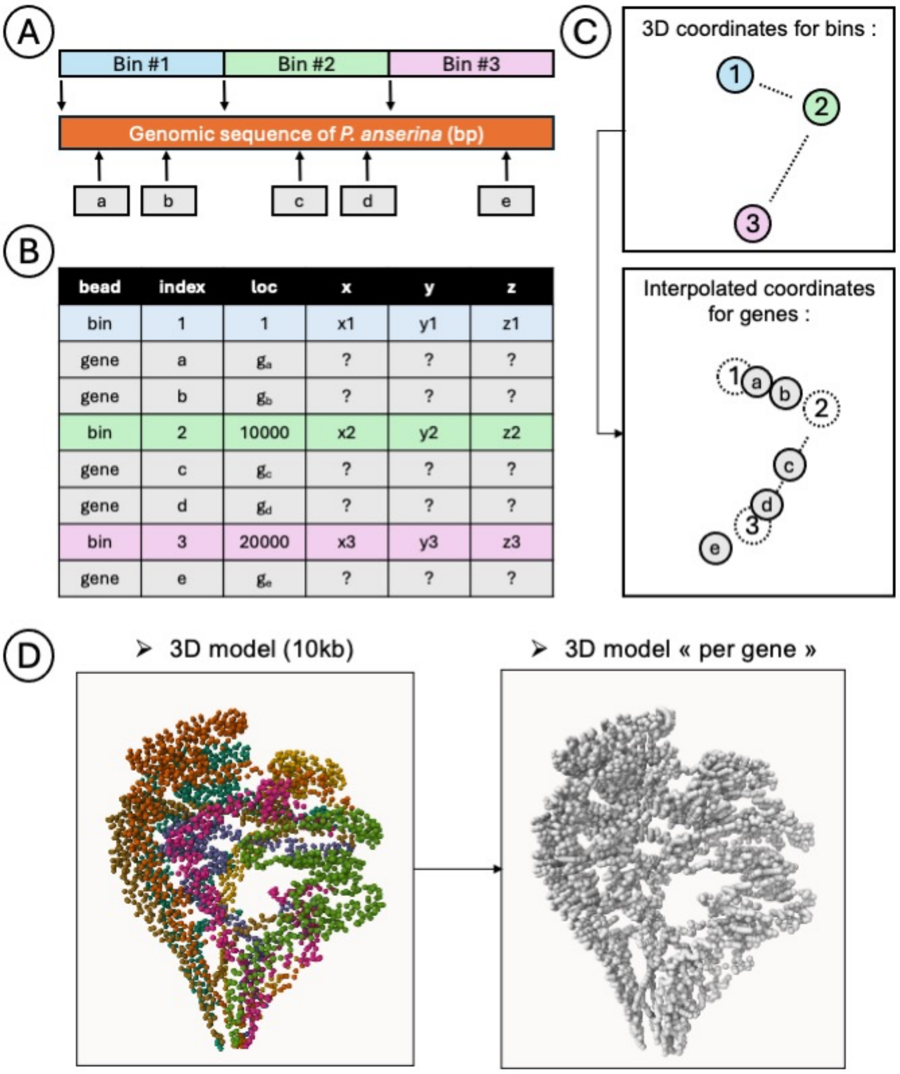


**Explanation of the procedure used to infer 3D coordinates of genes from the 3D coordinates for bins, as calculated with 3DGB software.** (**A**) The genomic sequence of *P. anserina* is used as a reference to map the starting positions of bins and the middle positions of genes, as described in an annotation file (see **DataFile 5**). (**B**) While the 3D coordinates of the bins are known, those of the genes are not. These are interpolated from those of the surrounding beads, as listed in a table containing all the coordinates (see **DataFile 5** for more details). (**C**) The 3D coordinates for genes *a* and *b* are derived from the 3D coordinates of beads #1 and #2, and the 3D coordinates for genes *c* and *d* are derived from the 3D coordinates of beads #2 and #3. (**D**) As a result, two models are available: the original "per bin" model, in which each bead represents a "bin" from the *P. anserina* genomic sequence, and the "per gene" model, in which each bead represents a gene based on its middle position in the genomic sequence.
