## Supplementary material for "Hi-C data from the filamentous fungus *Podospora anserina* and associated 3D models to visualize the spatial organization of its chromosomes": DataFile5: script_interpol.py

```
"""Interpolate genes from a PDB file containing a 3D genome structure.

It requires:
- a PDB file containing the genome structure,
- a fasta file containing the genome sequence,
- a csv file containing the genes positions,
- a resolution.

Author: Thibault Poinsignon
"""

import argparse
from mimetypes import suffix_map

import pandas as pd
import numpy as np     

from Bio import SeqIO
from biopandas.pdb import PandasPdb


def get_cli_arguments():
    """Command line argument parser.

    Returns
    -------
    argparse.Namespace
        Object containing arguments
    """
    parser = argparse.ArgumentParser()
    parser.add_argument(
        "--pdb",
        action="store",
        type=str,
        help="PDB file containing the 3D structure of the genome",
        required=True,
    )
    parser.add_argument(
        "--fasta",
        action="store",
        type=str,
        help="Fasta file containing the sequence of the genome",
        required=True,
    )
    parser.add_argument(
        "--resolution",
        action="store",
        type=int,
        help="HiC resolution from which the structure has been generated.",
        required=True,
    )
    parser.add_argument(
        "--annotation",
        action="store",
        type=str,
        help="BedGraph file containing the annotation of genes",
        required=True,
    )
    parser.add_argument(
        "--output",
        action="store",
        type=str,
        help="Output mmCIF file containing the annotated 3D structure of the genome",
        required=True,
    )
    return parser.parse_args()

def extract_chromosome_length(fasta_name):
    """Extract chromosome length from a FASTA file.
    
    Parameters
    ----------
    fasta_name : str
        Name of Fasta file containing the sequence of the genome
    
    Returns
    -------
    list
        List of chromosome lengths
    """
    chromosome_length_lst = []
    with open(fasta_name, "r") as fasta_file:
        print(f"Reading {fasta_name}")
        for record in SeqIO.parse(fasta_file, "fasta"):
            length = len(record.seq)
            print(f"Found chromosome {record.id} with {length} bases")
            chromosome_length_lst.append(length)
    return chromosome_length_lst

def get_base_pair_coordinates(atoms, chromosome_lengths, HiC_resolution):
    """Get base pair coordinate of structure's atoms.
    
    Parameters
    ----------
    pdb_name_in : str
        PDB file containing the 3D structure of the genome
    chromosome_length : list
        List with chromosome lengths
    HiC_resolution : int
        HiC resolution
    """
    atoms_bp_coor = pd.Series(dtype="int")
    for residue_number in atoms["residue_number"].unique():
        atoms_bp_coor_chromosome = pd.Series(range(0, chromosome_lengths[residue_number-1], HiC_resolution))
        atoms_bp_coor = pd.concat([atoms_bp_coor, atoms_bp_coor_chromosome])
    
    # Get the index of the beads without the ones already removed from the pdb file (by interpolate_missing_coordinates.py )
    atom_number = list(atoms["atom_number"]-1)
    # Delete the base pair coordinates corresponding to removed beads
    atoms_bp_coor.reset_index(drop=True, inplace=True)
    atoms_bp_coor = atoms_bp_coor.iloc[atom_number]
    atoms_bp_coor.reset_index(drop=True, inplace=True)
    
    atoms["bp_coordinate"] = atoms_bp_coor
    return atoms

def write_mmCIF(genes_output, mmCIF_name_out):
    out = open(mmCIF_name_out, "w")
    out.write("#\nloop_\n_atom_site.group_PDB\n_atom_site.label_seq_id\n_atom_site.label_atom_id\n_atom_site.label_asym_id\n_atom_site.label_comp_id\n_atom_site.Cartn_x\n_atom_site.Cartn_y\n_atom_site.Cartn_z\n_atom_site.B_iso_or_equiv\n")
    genes_output_str = genes_output.to_string(header=False, index=False, columns=["record_name", "atom_number", "residue_name", "chain_id", "atom_name", "x_coord", "y_coord", "z_coord", "b_factor"])
    out.write(genes_output_str)
    out.close()

def interpolate_genes(pdb_name_in, chromosome_lengths, annotation_name_in, HiC_resolution, mmCIF_name_out):
    """Interpolate genes according to a PDB file containing a 3D genome structure.

    Note:
    - The PDB file produced by Pastis is not readable by Biopython
    because the residue number column is missing.
    - We use instead the biopandas library. http://rasbt.github.io/biopandas/

    Parameters
    ----------
    pdb_name_in : str
        PDB file containing the 3D structure of the genome
    chromosome_length : list
        List with chromosome lengths
    annotation_name_in : str
        BedGraph file containing the annotation of genes
    HiC_resolution : int
        HiC resolution
    mmCIF_name_out : str
        Output mmCIF file containing the annotated 3D structure of the genome
    """
    pdb = PandasPdb().read_pdb(pdb_name_in)
    print(f"Number of beads read from structure: {pdb.df['ATOM'].shape[0]}")
    atoms = pdb.df["ATOM"]
    atoms = get_base_pair_coordinates(atoms, chromosome_lengths, HiC_resolution)
    atoms_output = pd.DataFrame(columns=atoms.columns)

    genes_bp_coordinates = pd.read_csv(annotation_name_in, sep="\t", header=None)
    genes_bp_coordinates.columns = ["residue_number", "bp_coordinate", "stop", "atom_name"]
    
    for residue_number in atoms["residue_number"].unique():
        atoms_chromosome = atoms[atoms["residue_number"]==residue_number]
        genes_bp_coordinates_chromosome = genes_bp_coordinates[genes_bp_coordinates["residue_number"]==residue_number]
        
        atoms_genes_chromosome = atoms_chromosome.merge(genes_bp_coordinates_chromosome, on="bp_coordinate", how="outer", sort=True, suffixes=("_x", None))
        atoms_genes_chromosome_interpolate = atoms_genes_chromosome.copy()
        atoms_genes_chromosome_interpolate["x_coord"] = atoms_genes_chromosome["x_coord"].interpolate(method='pchip', axis=0, limit_area='inside')
        atoms_genes_chromosome_interpolate["y_coord"] = atoms_genes_chromosome["y_coord"].interpolate(method='pchip', axis=0, limit_area='inside')
        atoms_genes_chromosome_interpolate["z_coord"] = atoms_genes_chromosome["z_coord"].interpolate(method='pchip', axis=0, limit_area='inside')
        chromosome = atoms_genes_chromosome_interpolate.loc[
        atoms_genes_chromosome_interpolate["record_name"] == "ATOM","residue_name"].iloc[0]
        atoms_genes_chromosome_interpolate["residue_name"] = chromosome
        chain_id = atoms_genes_chromosome_interpolate.loc[
        atoms_genes_chromosome_interpolate["record_name"] == "ATOM",
        "chain_id"].iloc[0]
        atoms_genes_chromosome_interpolate["chain_id"] = chain_id
        deleted_atoms = atoms_genes_chromosome_interpolate["x_coord"].isna().sum()
        print(f"Removed {deleted_atoms} genes from chromosome {residue_number} extremities")
        
        atoms_output = pd.concat([atoms_output, atoms_genes_chromosome_interpolate])
    
    atoms_output = atoms_output[atoms_output["record_name"]!="ATOM"]
    genes_output = atoms_output.dropna(subset=["x_coord"])
    
    genes_output.reset_index(drop=True, inplace=True)
    genes_output["residue_number"] = genes_output["residue_number"].astype(int)

    genes_output = genes_output.assign(record_name="ATOM",
                                        atom_number=list(range(1,len(genes_output)+1)),
                                        blank_1="",
                                        alt_loc="",
                                        blank_2="",
                                        insertion="",
                                        blank_3="",
                                        occupancy=1,
                                        b_factor=75,
                                        blank_4="",
                                        segment_id="",
                                        element_symbol="",
                                        charge=np.nan,
                                        line_idx=genes_output.index)
    genes_output.drop(["bp_coordinate", "atom_name_x",  "residue_number_x", "stop"], axis= 1, inplace=True)

    write_mmCIF(genes_output, mmCIF_name_out)
    print(f"Wrote {mmCIF_name_out}")
    print("Genes kept after interpolation:", len(genes_output))

    print("Genes loaded:", len(genes_bp_coordinates))
    print("Chromosomes in structure:", set(atoms["residue_number"]))
    print("Chromosomes in annotation:", set(genes_bp_coordinates["residue_number"]))
    print("bp range:", genes_bp_coordinates["bp_coordinate"].min(), "to", genes_bp_coordinates["bp_coordinate"].max())


if __name__ == "__main__":
    ARGS = get_cli_arguments()

    # Read Fasta file and get chromosome length
    CHROMOSOME_LENGTH = extract_chromosome_length(ARGS.fasta)

    # Assign chromosome number
    interpolate_genes(ARGS.pdb, CHROMOSOME_LENGTH, ARGS.annotation, ARGS.resolution, ARGS.output)
```
