## Supplementary material for "Hi-C data from the filamentous fungus *Podospora anserina* and associated 3D models to visualize the spatial organization of its chromosomes": DataFile7

### DataFile 7

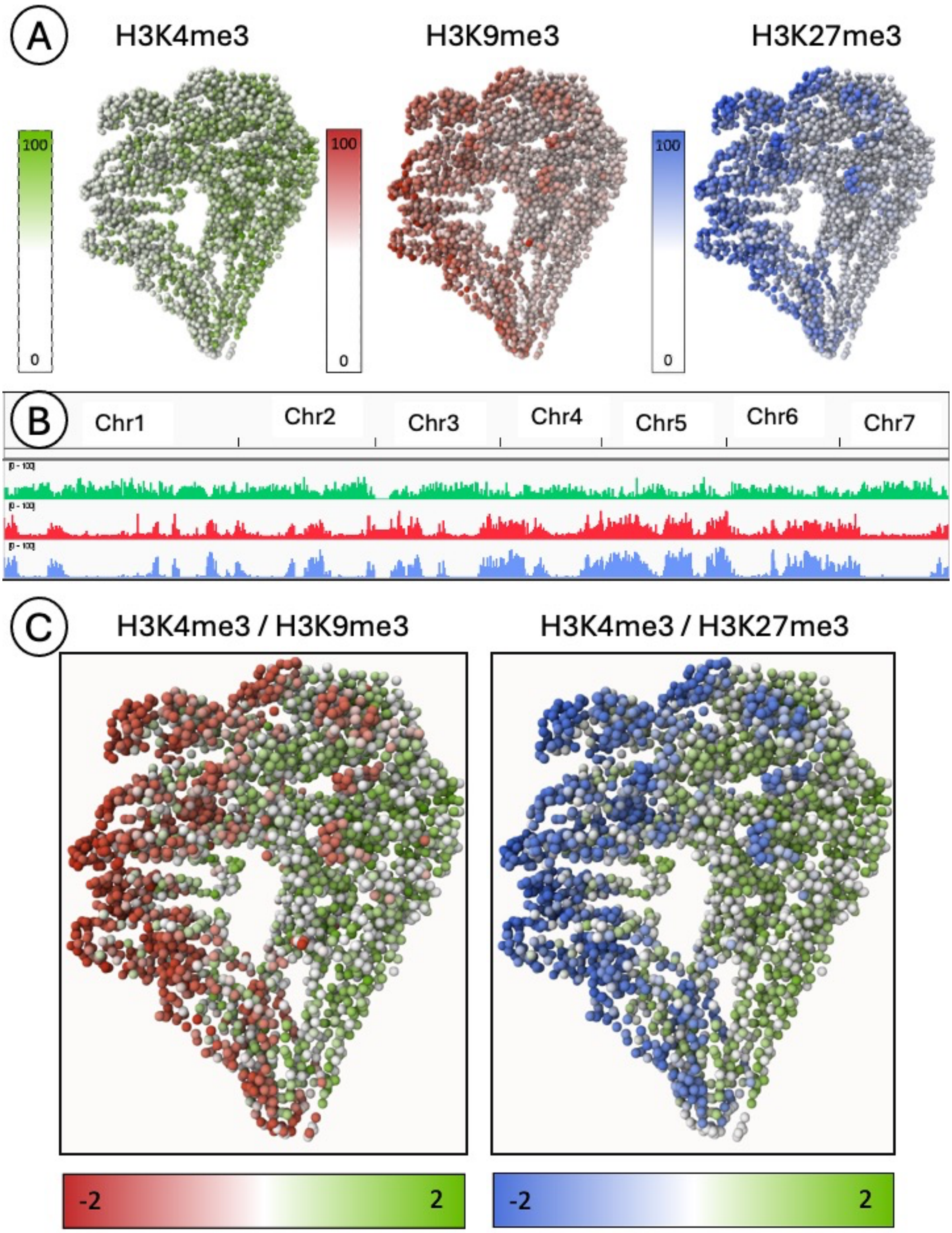


**Genome-wide patterns of three post-translational histone modifications, relating to open euchromatin (H3K4me3) and compact heterochromatin (H3K9me3 and H3K27me3).** The ChIP-seq data were collected from (1) and analysed using the pipeline available https://src.koda.cnrs.fr/AnakimGualdoni/3dgb-chipseq-analysis. (**A**) ChIP-seq signal intensities were calculated for each bead in the 3D model ('per bin') and used to colour the beads according to relative colour scales ranging from 0 (no signal) to 100 (very strong signal). (**B**) A linear representation of the same ChIP-seq signals is shown using a standard genome browser. Green, red and blue correspond to the histone marks H3K4me3, H3K9me3 and H3K27me3, respectively. (**C**) Representation of the relative intensities of the H3K4me3 and H3K9me3 or H3K27me3 signals is shown here. Positive values correspond to H3K4me3-specific regions (green), while negative values correspond to H3K9me3-specific (red) or H3K27me3-specific (blue) regions.
