## Supplementary material for "Hi-C data from the filamentous fungus *Podospora anserina* and associated 3D models to visualize the spatial organization of its chromosomes": DataFile8

### DataFile 8

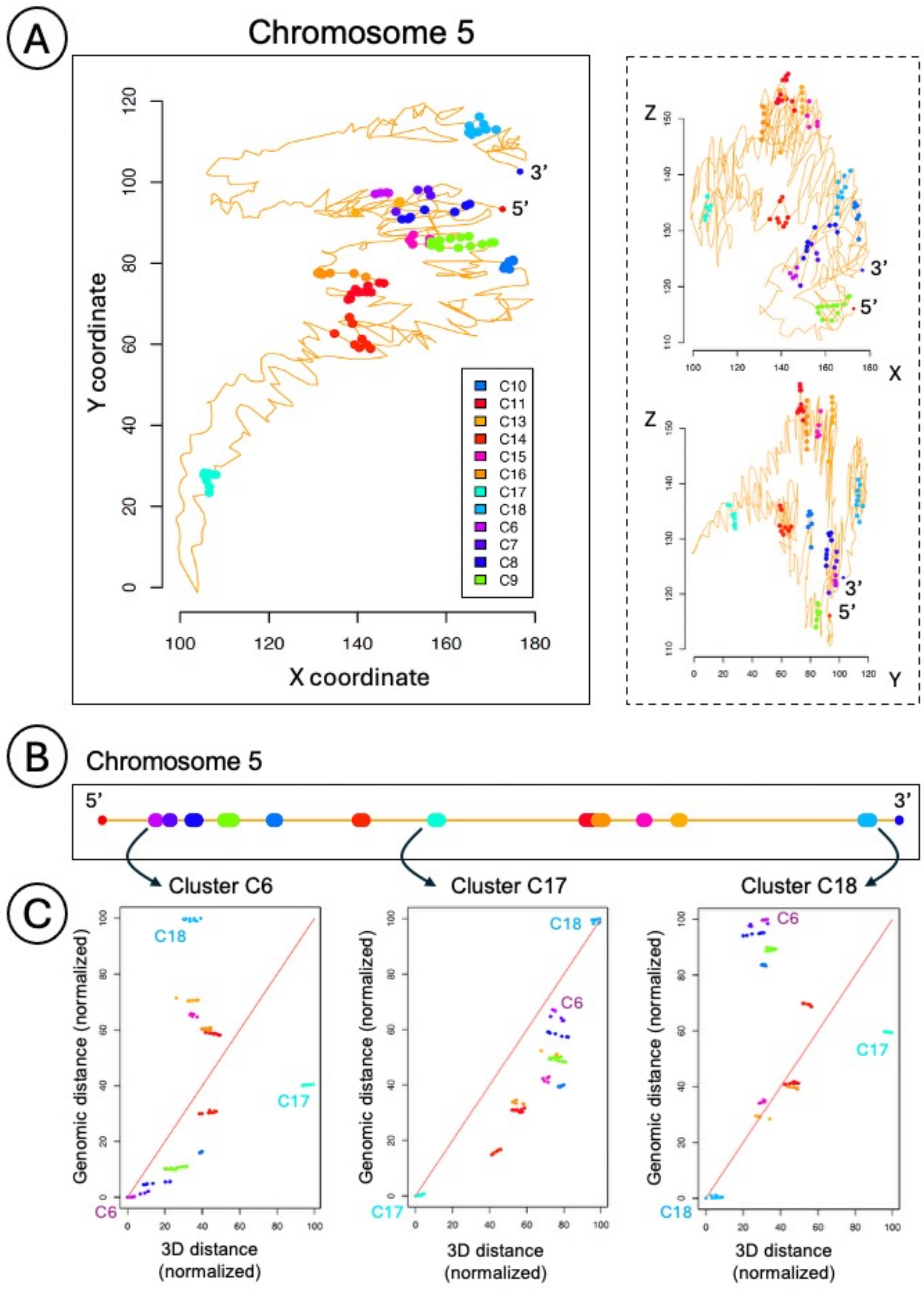


**Spatial localization of genes belonging to secondary metabolite clusters on chromosome 5 of *Podospora anserina*.** (**A**) Pairwise views of the 3D coordinates of genes. 12 clusters are known to be located on chromosome 5 (as described in (1)). They are shown here with different colors. Note that the 3D model used here is the 'per gene' model, meaning that each X, Y and Z coordinate represents the location of genes (see **DataFile4** for more explanations). (**B**) Linear representation of chromosome 5. The secondary metabolite clusters are represented with the same colors used in (**A**). Three genes belonging, respectively, to clusters C6, C17 and C18 were chosen as points of view to compare the genomic distances (in bp) with the Euclidean distance in 3D. Normalized values of the calculated distances are shown in (**C**).
